# Type IV pili increase phage encounter in marine cyanobacteria

**DOI:** 10.1101/2025.09.20.677514

**Authors:** Nieves M. Navarrete-López, Álvaro Sánchez-Carabantes, Jose M. Haro-Moreno, Mario López-Pérez, Maria del Mar Aguiló-Ferretjans, Joseph A. Christie-Oleza

## Abstract

Type IV pili are extracellular bacterial appendages with diverse functions, including biofilm formation, motility, conjugation, transformation, pathogenicity, and secretion. In planktonic marine cyanobacteria, they also enhance buoyancy and protect against grazing, yet their consequences for the oceanic carbon pump and microbial loop remain poorly understood. Pili production in dilute marine ecosystems entails ecological trade-offs, including high nutrient investment and, as we show here, increased vulnerability to phage infection. Using high-resolution electron microscopy, we observed cyanophages attaching along the entire length of extracellular pili, indicating that these filaments act as structural amplifiers of viral encounter in diffusion-limited environments. While piliated and non-piliated *Synechococcus* exhibited similar mortality in isolation, competition assays revealed a strong selective disadvantage for piliated cells, which were more rapidly depleted in mixed populations. This effect was most pronounced for cyanophage S-CAM7, which displayed higher pili-binding affinity and drove competitive exclusion of piliated hosts. To extend these findings to natural settings, we analysed single-cell amplified genomes (SAGs) observing pili genes encoded by 14.7% of total cyanobacterial SAGs. This proportion increased to 28.1% in virally infected SAGs (virocells; representing 3.8% of the total cyanobacterial SAGs), consistent with encounter-driven susceptibility of piliated cells in the environment. Together, our results identify type IV pili as double-edged structures: while conferring ecological advantages, they expose cells to viral attack by increasing encounter probability when pilus-attached phage are transport to the cell surface during pili retraction. This fitness cost may help explain the patchy distribution of pili in natural populations and the extreme sequence variability of pilins, reflecting an ongoing evolutionary arms race with phage.

## INTRODUCTION

Viruses are the most abundant biological entities in the ocean, estimated to exceed 10^30^ particles globally [1]. Through infection and lysis of microbial hosts, they profoundly influence marine ecosystems, driving nutrient turnover, modulating microbial diversity, and shaping evolutionary trajectories [2, 3]. Among their most important hosts are the picocyanobacteria *Prochlorococcus* and *Synechococcus*, which together dominate phytoplankton biomass in oligotrophic oceans and contribute up to one-third of global ocean primary production [4, 5]. Viral infection of these lineages exerts strong top-down control, with estimates that 10–40% of daily cyanobacterial mortality is attributable to phage lysis [6], and more recent studies suggesting 1-6% of picocyanobacterial cells being infected at any given time [7, 8]. Such mortality influences nutrient regeneration, carbon export, and the structure of microbial communities [9, 10]. Yet, despite this central role in shaping ocean ecosystems, the microscale determinants of phage–host encounters, the very first step of infection, remain poorly understood [11, 12]. Classical models of viral infection assume that the probability of successful infection scales with host density and phage abundance, modulated by adsorption rate constants derived from collision theory [11, 12]. Yet, field observations consistently reveal wide variability in infection prevalence, ranging from <0.01% to >40% of picocyanobacterial populations [8, 13, 14]. Such heterogeneity suggests that encounter probability is not governed by abundance alone, but also by structural and behavioural traits of the host that shape its effective visibility to viruses [11, 15-17]. Identifying these traits is central to understanding when and how cyanophages succeed in infecting their hosts.

One such trait is the presence of type IV pili [18, 19]. These extracellular appendages, assembled from pilin subunits, are widespread in bacteria and archaea and mediate diverse processes including motility, adhesion, biofilm formation and DNA uptake [20, 21]. In pathogenic bacteria, pili have been reported as phage receptors. For example, multiple *Pseudomonas aeruginosa* phages irreversibly adsorb to type IV pili, initiating infection when pilus retraction brings virions to the cell surface [18, 19]. In these systems, pili function as obligate gateways to infection, conferring susceptibility by providing a physical docking site.

In contrast, the ecological role of pili in marine picocyanobacteria remains less clear. Picocyanobacteria produce numerous long pili of ~6 nm diameter and up to ~10 µm in length [22]. While these structures have been linked to buoyancy regulation and grazing evasion [22, 23], their role in phage interactions in the marine realm has not been systematically investigated. Unlike pathogenic systems, where pili can function as essential receptors, cyanophages infecting *Synechococcus* and *Prochlorococcus* are believed to use alternative primary receptors, such as outer membrane proteins or lipopolysaccharides [24]. This distinction led us to hypothesize that, in marine picocyanobacteria, pili do not serve as strict entry points for infection, but instead act as modifiers of encounter probability. By extending the cell’s spatial footprint into the surrounding medium, pili may amplify the likelihood of viral contact and thereby shape infection risk.

From a physical perspective, pili dramatically extend the effective target radius of the cell (**Supplementary Fig. S1**). In diffusion-limited environments such as the open ocean, where viral motion is dominated by Brownian trajectories, even modest increases in encounter volume can substantially increase adsorption rates [17, 25]. A piliated cell, extending tens of microns into the surrounding medium [22], may therefore intercept more diffusing phage particles than an otherwise identical non-piliated cell. Importantly, not every encounter results in infection: phage adsorption efficiency depends on molecular compatibility and irreversible binding [26, 27]. Nonetheless, traits that amplify contact probability can bias infection risk, even in the absence of obligate receptors [28].

Whether pili actually make marine cyanobacteria more vulnerable to viruses has never been tested. Most experimental work on cyanophage–host interactions has focused on genomic characterization, adsorption assays, or bulk lysis dynamics [6, 29-32]. How structural appendages such as pili shape infection dynamics in mixed populations remains unexplored. At the same time, data indicating that piliated picocyanobacteria typically represent <25% of natural populations [22], suggest that pili production is constrained rather than universal. The forces maintaining this lower pilus-encoding frequency despite the ecological advantages they provide (i.e. avoid sinking and evade grazing predation), and whether viral pressure plays a role, remain unresolved. Here, we address this gap by providing the first experimental evidence that pili amplify virus–host encounter probability, imposing a hidden fitness cost under phage pressure. Combining transmission electron microscopy, monoculture and co-culture infection experiments, and single-cell genomic data, we test the hypothesis that pili increase viral encounters by expanding the effective contact radius of cells.

## MATERIALS AND METHODS

### Strains and culture conditions

*Synechococcus* sp. WH7803 and the non-piliated knockout mutant (Δpili) were grown in ASW medium [33]. Δpili strain was generated previously in our lab by replacing the *pilA1* (SynWH7803_1795) and *pilE* (SynWH7803_1796) genes with a gentamycin resistance cassette via a double homologous recombination strategy [22]. The mutant is fully segregated, axenic, and exhibits a stable sinking phenotype (details in [22]). Experiments were performed using 30 ml cultures contained in 25 cm^2^ rectangular cell culture flasks (Falcon) with vented caps. Cultures were incubated under optimal growth conditions i.e. at 22 °C at a light intensity of 10 μmol photons m^−2^ s^−1^ with orbital shaking (160 r.p.m.). Cyanobacterial culture growth was routinely monitored by flow cytometry (BD Accuri, BD Biosciences) operated with BD FACSAccuri software.

### Cyanophage propagation and plaque assays

Characterised cyanophages S-CAM7, S-PM2 and S-RSM4 [29, 34, 35] were propagated on *Synechococcus* sp. WH7803 grown in 250 mL acid-washed Erlenmeyer flasks containing 100 ml of ASW under optimal growth conditions. Following complete lysis, lysates were adjusted to 6% NaCl and treated with chloroform (10% w/v) to detach viral particles from cell debris. The aqueous fraction was recovered, centrifuged (4000 x g, 15 min) and supernatant further filtered through 0.22 μm PES syringe filters (Corning). Phage filtrates were concentrated ~100x using Amicon Ultra filter columns (100 kDa MWCO; Merck). Phage concentrate was filter-sterilised through 0.22 μm PES syringe filters and stored in the dark at 4 ºC until further use. Phage titers were determined by dilution-to-extinction in 96-well microplates overlayed by *Synechococcus* sp. WH7803 and assessed by culture lysis after a 7-days incubation under optimal conditions.

To both confirm phage titers and evaluate the fitness cost associated with piliated *versus* non-piliated strains, plaque assays were carried out as previously described [36]. Serial dilutions of phage stocks were plated with either *Synechococcus* sp. WH7803 or Δpili, and incubated under continuous light (10 μmol photons m^−2^ s^−1^) until plaques became visible. For size comparison, only well-separated plaques were analysed to ensure accurate measurement. Plaque diameters were determined using ImageJ [37], with pixel-to millimetre conversion based on a reference image containing a 1 mm scale bar corresponding to 25 pixels.

### Transmission electron microscopy

A volume of 100 μL obtained from an optimally-grown *Synechococcus* sp. WH7803 culture (1 × 10^8^ cells ml^−1^) was mixed with 10 μL of cyanophage concentrate (10^10^ plaque forming units mL^−1^; multiplicity of infection, MOI=10) and incubated in the dark at 4 °C for 30 minutes to allow phage adsorption. Preliminary tests with different MOIs indicated that 10 was the most reproducible ratio for assessing pili–virus interactions. Subsequently, 10 μL of the mixture were gently applied onto a glow-discharged formvar/carbon-coated copper grid and incubated for 10 minutes to allow cell attachment. Excess liquid was carefully blotted off, and negative staining was performed by applying a drop of 2% (w/v) uranyl acetate for 5 minutes. The stain was then blotted off, followed by a gentle rinse with Milli-Q water for 5 minutes. After blotting the excess, grids were imaged using a high resolution electron microscope Talos F200i (TheromScientific) JEOL 2011 transmission electron microscope equipped with a CETA acquisition camera (VELOX). Because no high-throughput method was available for quantifying phage bound to pili, transmission electron micrographs were randomly acquired and analysed by two operators using defined morphological criteria. The cyanophages S-CAM7, S-RSM4, and S-PM2 are T4-like viruses with contractile tails and well-defined baseplates. A phage was scored as “bound to pili” when its base structure appeared distinctly locked-in to a *Synechococcus* pilus, with tail fibers tightly engaged and increased electron density at the point of contact. Only viruses with intact capsids were counted.

### Infection assays

Infection assays with each one of the three phage types –S-CAM7, S-RSM4, and S-PM2– were performed in mono- and co-cultures of piliated and non-piliated *Synechococcus* sp. WH7803 genotypes. Dilute *Synechococcus* cultures (~10^6^ cells ml^−1^) that would then allow downstream analyses, and a low phage load was used (~300 active phage particles per 30 ml culture; corresponding to an MOI of 1 × 10^−5^) in order to amplify phage-host encounter time differences over multiple infection cycles. After infection, cultures were incubated at 22 °C at a light intensity of 10 μmol photons m^−2^ s^−1^ with orbital shaking (160 r.p.m.) to avoid Δpili mutant sinking. Cultures were sampled daily over 8 days, taking 100 μL for flow cytometry, and 1 mL for cell pelleting by centrifugation (14,000 x g for 1 min) and stored at –20 °C until further processing.

### DNA extractions and qPCR determination of piliated and non-piliated cells in co-cultures

DNA was extracted from cells pelleted from 1 ml culture samples using an optimized protocol. Pellets were resuspended in 200 μl Milli-Q water, and ~0.1 g of glass beads (0.1 mm in diameter) was added, and vortexed vigorously. Then, samples underwent three cycles of heating (96 °C, 3 min) followed by 3 min of bead beating (Disruptor Genie from Scientific Industries). After processing, suspensions were centrifuged (14,000 × g, 10 min), and the supernatant was used directly as the DNA template for qPCR. Strain-specific quantification of piliated and non-piliated *Synechococcus* sp. WH7803 was performed by qPCR using two primer pairs: the WT strain was detected with primers targeting *pilA*, and the Δpili mutant with primers targeting the gentamycin resistance cassette. Primer sequences are listed in **Supplementary Table S1**. Primers were designed using Primer-BLAST (NCBI) and experimentally optimized to achieve equivalent amplification efficiencies across primer pairs (e.g., 97.51% efficiency with 1 μM of the *pilA* primer set and 97.62% efficiency with 500 nM of the gentamycin primer set). Controlled mixtures of WT and Δpili cells at ratios of 10:1, 1:1, and 1:10 confirmed accurate quantification of both genotypes, with reliable differentiation maintained above 10^6^ cell pellets.

qPCR reactions (10 μl) contained 5 μl PowerUp SYBR Green Master Mix (Applied Biosystems), 500 nM of the gentamycin primer pair or 1 μM of the *pilA* primer pair, 1 μl template DNA, and nuclease-free water. Each plate included a dilution series in triplicate for standard curve generation. Cycling conditions were 50 °C for 2 min, 95 °C for 10 min, and 40 cycles of 95 °C for 15 s and 60 °C for 1 min. Amplifications were run on a QuantStudio 3 Real-Time PCR System (Applied Biosystems). All reactions were performed in triplicate. Raw Ct values were exported for analysis and normalized against flow cytometry-based cell counts.

### Single-Cell Genome analysis

Available single-cell amplified genomes (SAGs) of marine picocyanobacteria [38, 39] were downloaded from the NCBI BioProjects PRJEB33281 and PRJNA445865, and checked for genome completeness and contamination using checkm2 v1.1.0 [40]. SAGs with ≥ 50% completeness and ≤5% contamination were selected, resulting in 690 *Prochoroccocus* and 31 *Synechococcus* SAGs (**Supplementary Table S2**). Actively phage-infected cells –excluding lysogenic infections– were identified amongst the SAGs using VIBRANT v1.2.1 [41], and confirmed either with geNomad v1.11.1 [42], Virsorter2 v2.2.4 [43], or both (**Supplementary Table S3**). Coding-DNA sequences from the 721 Cyanobacterial SAGs were predicted using prodigal v.2.6.3 [44], and proteins screened using hmmscan v3.4 [45] against the full KEGG dataset (downloaded in November 2024, [46]) for hits matching the structural pili genes *pilB* (K02652) and *pilC* (K02653) as well as single-copy house-keeping genes *recA* (K03553) and *rpoB* (K03043) (**Supplementary Table S4**). For each protein, only the best hit was considered, and hits above the recommended trusted scores were kept. The proportion of piliated cells amongst the total collection of SAG and subset of virocells was calculated by normalising the average number of *pilB* and *pilC* by the average number of house-keeping genes *recA* and *rpoB*.

## Data analysis

All statistical analyses were performed in R [47] using RStudio [48]. Unless otherwise indicated, values represent the mean ± standard deviation (SD) of three independent biological replicates. Differences between treatments were assessed using Welch’s two-sample *t*-tests, restricted to days 3–8 post-infection. Resulting *p*-values were corrected for multiple comparisons using the Benjamini–Hochberg false discovery rate (FDR) method, and only FDR-adjusted values < 0.05 were considered significant.

## RESULTS AND DISCUSSION

### Cyanophage bind to type IV pili of *Synechococcus* sp. WH7803

We exposed *Synechococcus* sp. WH7803 to three well-characterized cyanophage: S-CAM7, S-PM2, and S-RSM4 [29, 34, 35]. These T4-like phage of cyanobacteria have extensively been referred to as cyanomyoviruses, owing to their contractile tails and genomic similarity to members of the *Myoviridae* family [30]. Using transmission electron microscopy (TEM) at an MOI of 10, we observed frequent virion attachment to the pili of *Synechococcus* (**Fig. 1A** and **Supplementary Fig. S2**). While such interactions are well documented in pathogenic bacteria [18, 19], they have not previously been reported in marine cyanobacteria. This result prompted a closer examination of phage–pili interactions in marine picocyanobacteria.

**Figure 1.**
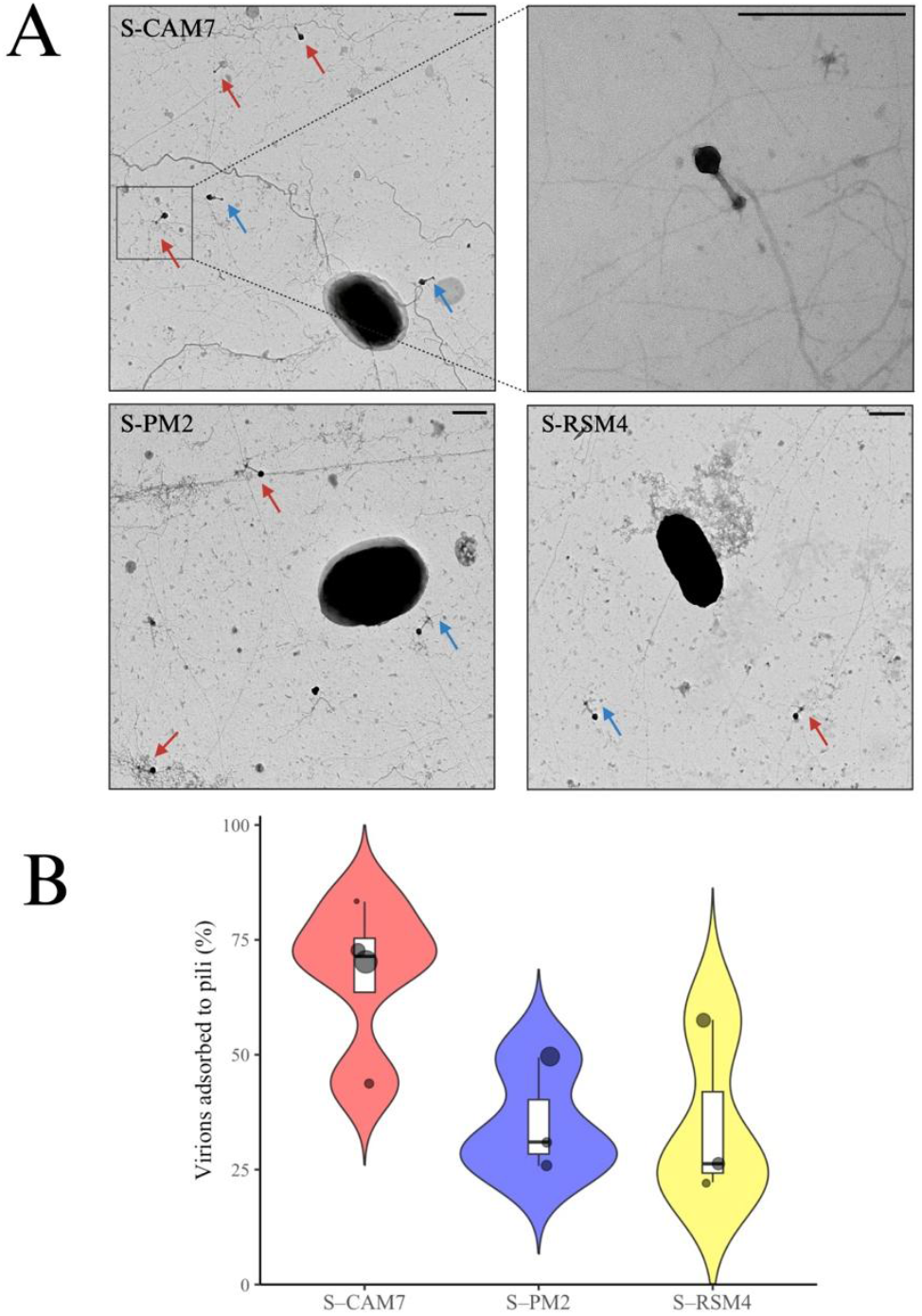
Cyanophage binding to type IV pili in *Synechococcus* sp. WH7803. **(A)** Transmission electron microscopy images of piliated *Synechococcus* sp. WH7803 in the presence of cyanophages S-CAM7, S-PM2, and S-RSM4. Red arrows indicate virions adsorbed to type IV pili, and blue arrows indicate those that were not adsorbed. Scale bars: 500 nm. **(B)** Quantification of phage-pili interactions. Violin plots show the distribution of adsorption frequencies based on visual counts of virions across multiple TEM fields, with embedded boxplots indicating medians and interquartile ranges (n = 225, 87, and 154 particles for S-CAM7, S-RSM4, and S-PM2, respectively).

High-resolution TEM revealed that phage attachment to type IV pili was not confined to distal tips, but occurred along the entire filament, including mid-pilus regions (**Fig. 1A**). Such positional variability has been described in other systems—for example, F-type phage adsorb specifically at *E. coli*’s pilus tip [49], whereas ϕ6 binds along the sides of the pilus of *Pseudomonas* [50]. In *Synechococcus*, cyanomyoviruses attached both distally and laterally, indicating that pili serve as multivalent scaffolds for docking. These observations led us to hypothesize that by extending the surface available for adsorption, pili enlarge the effective encounter cross-section of the host, thereby increasing the probability of viral contact in diffusion-limited environments.

All three cyanophages analysed, S-CAM7, S-RSM4, and S-PM2, are T4-like viruses with contractile tails and well-defined baseplates. When bound to pili, their base structures appeared distinctly locked-in, with tail fibers tightly engaged and increased electron density at the contact point (**Supplementary Fig. S2**), consistent with an attachment-ready state [51]. Binding frequencies, however, differed markedly between phages: S-CAM7 displayed consistently higher pili-association, with a median of 70% and relatively tight distribution, whereas S-PM2 and S-RSM4 showed lower median values (43% and 44%, respectively), and broader and more heterogeneous distributions (**Fig. 1B**). These contrasting profiles suggest differences in receptor compatibility, likely underlying receptor affinity or divergence.

These observations suggest that type IV pili in *Synechococcus* support viral docking, facilitating virion-cell surface encounter during pilus retraction. Unlike Siphoviridae, which lack tail fibers and rely solely on baseplate proteins [52], the cyanomyoviruses studied here possess contractile tails and specialized fibers, likely enabling stronger or more stable pili interactions. Although we did not characterize tail proteins directly, the distinct baseplate engagement observed in TEM images is consistent with a model in which either fibers or baseplate domains mediate pilus recognition [53]. Together, these findings establish a structural basis for our hypothesis that pili amplify host–virus encounter probability in marine cyanobacteria.

### Type IV pili enhance phage encounter in marine cyanobacteria

In the surface ocean, viruses typically outnumber their microbial hosts by one to two orders of magnitude [12]. Yet, despite this overwhelming numerical advantage, infection rates in picocyanobacteria remain strikingly variable, ranging from <0.005% [13] to nearly half of the population [14], with recent studies suggesting <6% of picocyanobacteria being infected at any given time [7, 8]. Such variability points to more than simple host and virus abundance. Instead, encounter efficiency is likely modulated by macroscale biogeographic factors [8] and, most importantly, microscale host traits that determine how visible a cell is to viruses [11, 14-17]

One such trait is the production of type IV pili. In contrast to *Synechocystis* sp. PCC6803, which produces both thick and thin pili [46], marine *Synechococcus* sp. WH7803 forms numerous filaments of uniform diameter (6 nm) that can extend up to 10 µm from the cell surface [22]. These extracellular structures dramatically enlarge the zone around the cell where first contact with phage might occur. In nutrient-poor ocean waters, where cells and viruses are highly dilute and move largely by diffusion, even modest increases in target size can translate into substantially higher encounter rates. Although not every collision leads to irreversible binding, simply raising contact probability increases the overall likelihood of successful infection [17, 26, 28].

Phage infection itself unfolds as a sequence of diffusive events: viral motion, collision with the host, reversible binding, and, if molecularly compatible, irreversible attachment [11, 17]. This process can be formalized by the phage adsorption constant (*k*), expressed as:

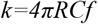

where *R* is the effective host radius, *C* is the phage diffusion coefficient, and *f* the probability of collision leading to attachment.

Here, we hypothesized that type IV pili act not as strict entry receptors, but as amplifiers of encounter probability by enlarging the effective radius of the host. This idea builds on Smoluchowski theory [25], applied to phages [26], which captures the key determinants of adsorption: how fast phage diffuse, how large the target is, and how likely contact results in binding [17]. Assuming *C* and *f* remain constant, our geometric modelling predicts that pili can expand the effective encounter radius of *Synechococcus* WH7803 by up to 20-fold, resulting in a proportional increase in *k*. This estimate arises directly from cell geometry: while non-piliated cells are 1 µm in diameter, pili can extend up to 10 µm in the surrounding medium [22]. By creating an extended, filamentous structure that intercepts diffusing particles, pili increase the likelihood of contact without requiring changes in molecular specificity. However, this 20-fold value represents a theoretical upper bound, assuming uniform pilus coverage and constant adsorption efficiency, as well as pili forming a 3D spherical structure around the cell. The true effect is likely smaller but still higher relative to non-piliated cells, and has not previously been tested in marine cyanobacteria (see derivation in **Supplementary Note 1**).

Patchy pili production in *Synechococcus* populations [54] may represent a strategy to balance ecological function with viral risk. Heterogeneous expression ensures that while a fraction of cells benefits from pili-mediated interactions with the environment, others remain less exposed to phage attack, providing a population-level buffer against infection. Such phenotypic diversity could stabilize populations under fluctuating viral pressure and maintain coexistence of piliated and non-piliated lineages. At the community scale, this heterogeneity influences the strength of phage–host interactions, potentially modulating turnover rates and shaping microbial community structure in the open ocean [27, 55]

Together, these considerations suggest that pili could substantially amplify the likelihood of virus– host encounters in the open ocean. Yet, whether this predicted increase in contact probability actually makes marine cyanobacteria more susceptible to phage has never been tested. To address this, we next performed monoculture infections of *Synechococcus* WH7803, directly comparing piliated and non-piliated strains to determine whether pili translate into higher phage-induced mortality.

### Piliated cells are more susceptible to phage than co-existing non-piliated counterparts

Although TEM revealed frequent phage attachment to type IV pili, bulk assays such as adsorption tests and plaque expansion proved inconclusive between wild type *Synechococcus* sp. WH7803 (WT, piliated) and the pili knockout mutant (Δpili; where genes *pilA1* and *pilE* were deleted; [22]). The fragility of pili means they shear during centrifugation, leaving attached phage in the supernatant during conventional phage-host attachment measurements. Similarly, measurements of plaque sizes yielded similar results between WT and Δpili strains (**Supplementary Fig. S3**).

We then tested liquid monoculture mortality of *Synechococcus* WH7803 (WT, piliated) and pili-deficient mutant (Δpili; [22]) when infected independently with three cyanophages (S-CAM7, S-PM2, and S-RSM4). Again, these monoculture experiments displayed nearly identical culture lysis dynamics, with no statistically significant differences across all virus–host combinations (Welch’s t-tests, FDR-corrected *p* > 0.05; **Fig. 2A-C**). Slightly greater WT depletion was observed late in S-CAM7 and S-PM2 infections (**Fig. 2A, C**), but the effect was small and non-significant. These results indicate that, in monoculture settings, where cells are in relatively high abundance and not in competition, pili do not significantly increase phage-induced time of mortality. At first glance this appears to contradict our encounter amplification hypothesis, however, in the absence of competition in pure culture settings, the time used for host-phage adsorption may be insignificant (within the minute range) compared to the overall infection cycle (i.e. phage burst time), this being over 8 h for these cyanophages [56].

**Figure 2.**
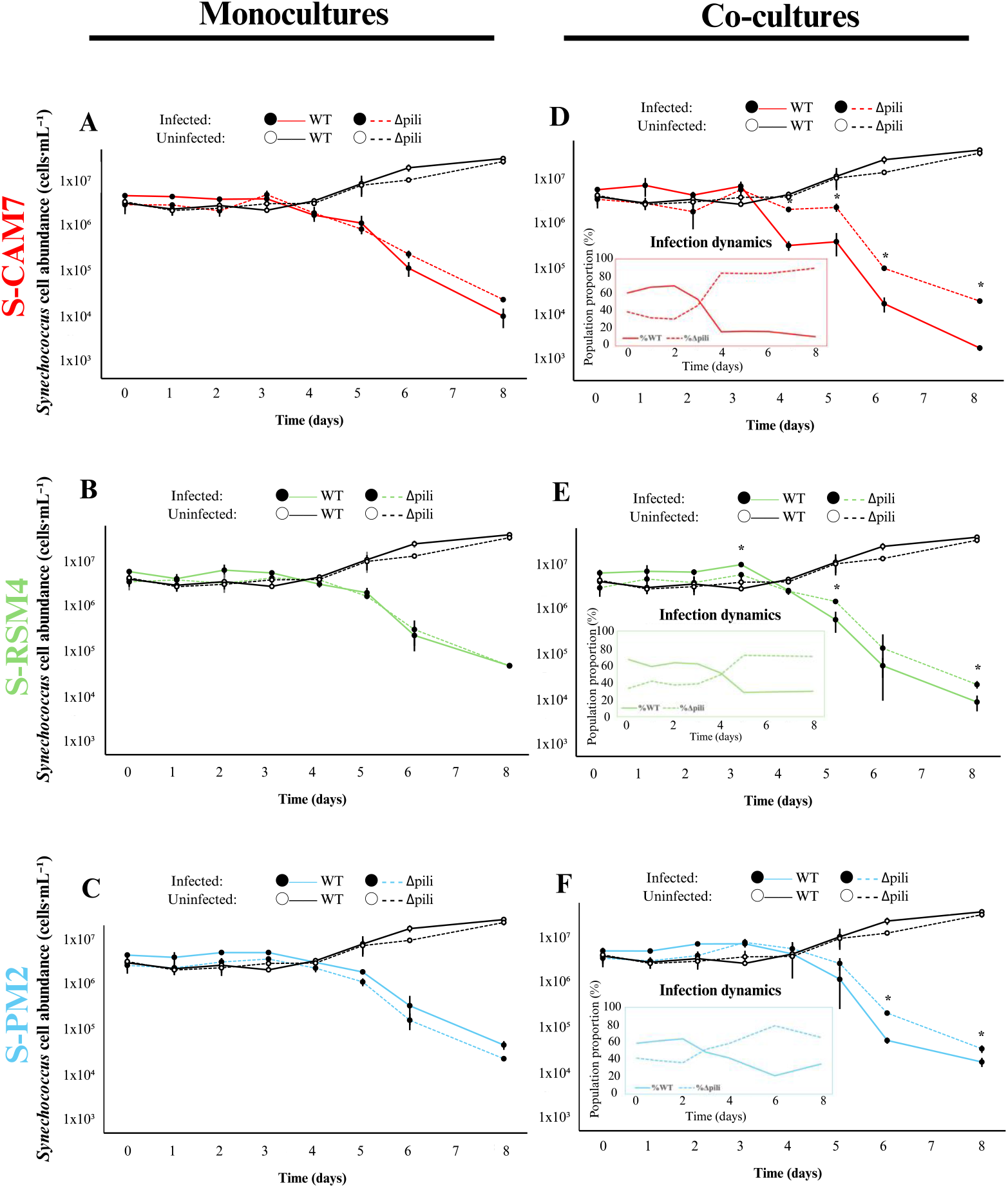
Phage infection dynamics in monocultures and co-cultures of piliated and non-piliated *Synechococcus* sp. WH7803. Piliated (WT) and non-piliated (Δpili) strains were either grown separately (monocultures, panels **A–C**) or together at a 1:1 ratio (co-cultures, panels **D–F**), and exposed to the cyanophages S-CAM7 (**A, D**), S-PM2 (**B, E**), and S-RSM4 (**C, F**) at a multiplicity of infection of 10^−5^. Population dynamics were monitored over 8 days by flow cytometry and strain-specific qPCR. Data represent mean ± SD from three independent biological replicates. Statistical significance was assessed using Welch’s t-test with false discovery rate (FDR) correction; asterisks denote days where WT and Δpili differed significantly (p < 0.05).

The contrast between theoretical expectation and monoculture outcome prompted us to test whether the cost of pili becomes evident under direct competition, i.e. when phage are amongst mixed piliated and non-piliated cells, where encounter differences can translate into relative fitness effects. We co-cultured WT and Δpili cells at a 1:1 ratio and exposed them again to the three cyanophages (S-CAM7, S-PM2, S-RSM4). Population dynamics were tracked over 8 days using flow cytometry and quantitatively differentiating both genotypes by strain-specific qPCR. Under these settings, piliated cells showed a significant disadvantage *versus* non-piliated *Synechococcus* in the presence of phage. In all viral treatments, piliated cells progressively declined in relative abundance, while Δpili cells became visibly dominant (**Fig. 2D-F**). This competitive shift was detectable from day 3 onwards and was most pronounced in S-CAM7 infections – the phage with the highest pili binding affinity (**Fig. 1B**). While WT to Δpili cell ratios in the co-cultures started at 60%:40%, during S-CAM7 infection, the proportion of piliated WT cells declined to 16.3% by day 4 and stabilised at 10.7% by day 8, while non-piliated Δpili cells became 89.3% of the population (**Fig. 2D**). Under S-PM2 and S-RSM4, similar but less pronounced shifts were observed, with piliated populations dropping to 31.44% and 28.35% of the co-culture by day 5, respectively, and a consequent increase in relative abundance of the non-piliated mutant (**Fig. 2E–F**).

Phage suppressed piliated cells under competitive conditions because pili increased host–virus encounter probability, converting a cryptic trait in monoculture into a clear disadvantage under selective settings. Curiously, our results mirror natural patterns: the strong selective disadvantage we observed aligns with the consistently low global frequency of piliated picocyanobacterial in the global oceans (i.e. <25% of cyanobacterial cells encode for pili in the ocean; [22]), and may reflect a virus-driven equilibrium. In dilute marine waters, where infection is encounter-limited, pili increase host visibility to phage, biasing adsorption events toward piliated cells and, possibly, suppressing their abundance. Viral lysis acting across cyanobacterial lineages could therefore maintain piliated cells at low levels, balancing the ecological benefits of pili, such as buoyancy and grazing resistance [22], against their cost under phage predation.

### Increased viral infections in pili-encoding picocyanobacteria in natural marine communities

To assess whether piliated cells are disproportionately targeted by phage in natural marine environments, we analysed single-cell genomic datasets [38, 39, 57] with >50% completeness (690 *Prochlorococcus* and 31 *Synechococcus* SAGs; **Supplementary Table S2**). The proportion of piliated cells amongst *Prochlorococcus* SAGs (12.7%) was lower than *Synechococcus* SAGs (59.4%), but within the ranges previously reported [22]. Approximately 3.8% of all *Prochlorococcus* and *Synechococcus* SAGs were classified as virocells (**Supplementary Table S3**), i.e. cells actively engaged in phage infection [58], a proportion within the expected number of infected cells in natural marine environments at any given time [7, 8]. Most interestingly, while the overall average of piliated picocyanobacterial SAGs was 14.7% (i.e. due to the high abundance of *Prochlorococcus* amongst the dataset), this percentage almost doubled to 28.1% when considering only the virocell SAGs (**Supplementary Table S4**). Overall, this suggests that piliated cells are much more infected in natural ecosystems and confirms the ecological burden of producing pili in the presence of viral predators.

Furthermore, the abundance of pili amongst the natural picocyanobacterial community (i.e. 14.7%) is strikingly close to the equilibrium reached in our piliated : non-piliated co-cultures under phage pressure, suggesting that viral selection may keep piliated cells as a persistent minority in natural populations. These patterns are consistent with a kill-the-winner dynamic [55], where traits that increase encounter probability (here, pili) are suppressed by viral predation in open, dilute marine environments. If infection were inevitable in all piliated cells, or pili production did not confer a strong ecological advantage, this trait would likely have been purged from cyanobacterial populations. Instead, its persistence despite a clear viral cost suggests that the ecological benefits of pili are sufficient to offset the risk of infection [6, 59]. Together, these results point to encounter-driven trade-offs under phage pressure as a key mechanism maintaining piliated hosts as a minority in natural cyanobacterial populations.

## CONCLUSIONS

Our study uncovers a previously unrecognized role for type IV pili in marine picocyanobacteria: they act as encounter amplifiers, enlarging the effective host radius and increasing viral encounter probability. This reveals an ecological trade-off for these, *a priori*, highly beneficial structures that help retain cellular positioning within the water column and protect against grazing. Hence, we prove that pili expose cells to elevated phage attack, linking these structures directly to microbial survival strategies and the oceanic viral shunt. We further show that, despite the benefits, viral fitness costs keep piliated picocyanobacterial cells in low abundance (~10-25%), suggesting patchy piliation as a potential bet-hedging strategy. Yet key questions remain: how pili expression is regulated in natural populations, whether structural diversity determines phage specificity, and to what extent pili-mediated encounters contribute to global viral lysis and carbon cycling. Addressing these gaps will clarify how a simple surface appendage shapes virus–host dynamics and global ecological processes in the open ocean.

## ACKNOWLEDGEMENTS

The authors thank Dr. Busquets Bisbal for the acquisition of the TEM images, and Prof. Scanlan and Dr Puxty at the University of Warwick for providing the cyanophage used in the study.

## AUTHOR CONTRIBUTIONS

**Nieves M. Navarrete-López**: Conceptualization, Methodology, Validation, Formal analysis, Investigation, Data curation, Writing – original draft, Visualization. **Álvaro Sánchez-Carabantes**: Methodology, Formal analysis, Investigation, Data curation, Writing – review & editing. **Jose M. Haro-Moreno**: Methodology, Formal analysis, Data curation, Writing – review & editing. **Mario López-Pérez**: Methodology, Validation, Formal analysis, Data curation, Writing – review & editing. **Maria del Mar Aguiló-Ferretjans**: Methodology, Investigation, Writing – review & editing. **Joseph A. Christie-Oleza**: Conceptualization, Validation, Data curation, Resources, Writing – original draft, Visualization, Supervision, Funding acquisition, Project administration.

## SUPPLEMENTARY MATERIAL

Supplementary material is available online.

## DATA AVAILABILITY

All data are available in the main text or the supplementary materials.

