## Supplementary Information (Suppl Table S1 and Figures S1-S3) for "Type IV pili increase phage encounter in marine cyanobacteria"

**This file includes**

**Supplementary Table 1** and **figures S1** to **S3.**

**Supplementary Note 1:** Smoluchowski adsorption model for piliated versus non-piliated cells.

**Supplementary Table 1.** Primers used for qPCR amplification during infection assays.

| Primer name | Sequence (5'-3') | Annealing temperature |
|---|---|---|
| **WT (*pilA* locus)** - F | TCACAGCTTTCACCGGATGG | Tm 60,32 ºC |
| **WT (*pilA* locus)** - R | GTCGAGGCTTGTAGTCGTGT | Tm 59,75 ºC |
| **Δpili (gentamicin locus)** - F | TTTCGGTCGTGAGTTCGGAG | Tm 60,04 ºC |
| **Δpili (gentamicin locus)** - R | GCAAGCGCGATGAATGTCTT | Tm 59,90 ºC |

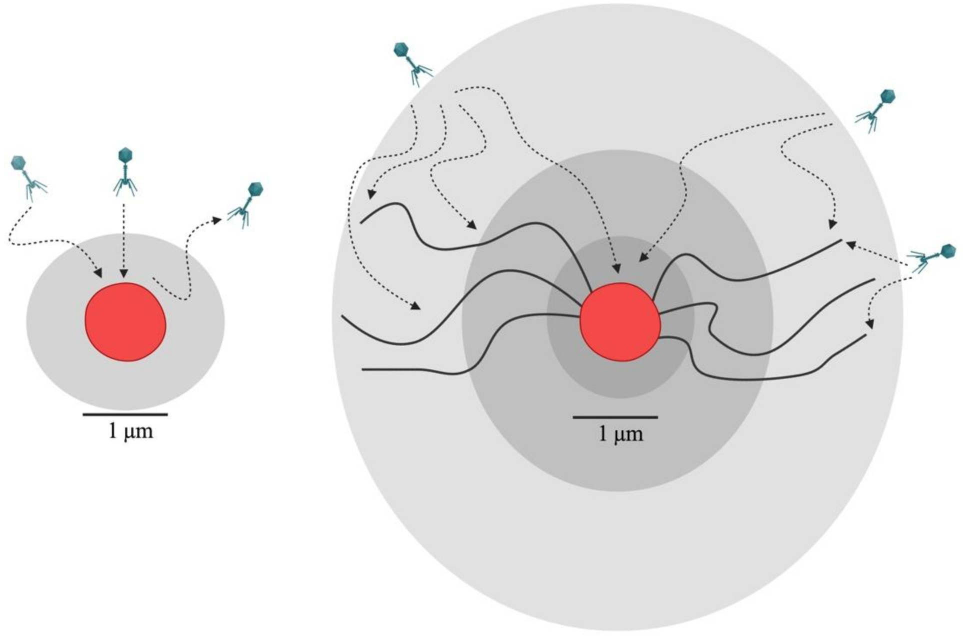


**Supplementary Figure S1. Type IV pili expand host encounter volume in marine picocyanobacteria.** Schematic representation of piliated (right) and non-piliated (left) cells. In piliated cells, filaments (10 µm long) extend up to 10 times the cell diameter and radiate from the membrane, creating a three-dimensional zone where viruses may first make contact. Dashed arrows illustrate potential phage trajectories, either striking the membrane directly or colliding first with a pilus. While infection ultimately requires surface attachment, pili increase the effective encounter volume and may facilitate viral delivery to the cell surface.

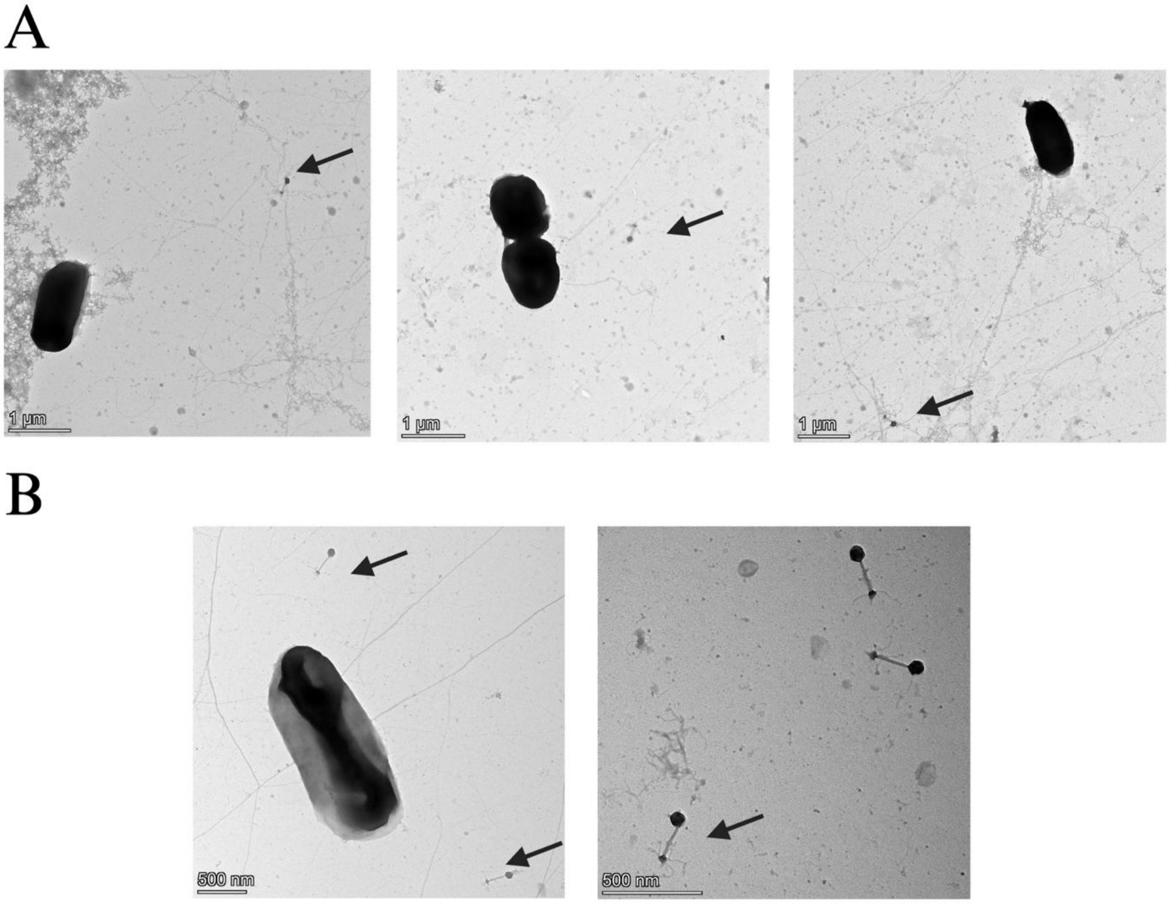


**Supplementary Figure S2. Representative TEM images illustrating classification criteria: bound virions with baseplate engaged on pili (A) *versus* free unbound virions (B).** (**A**) S-RSM4 absorbed to pili. Virions were scored as bound when their baseplate was directly locked onto a pilus extending from a host cell, with tail fibers tightly wrapped around the pilus and increased electron density at the contact point (black arrows). (**B**) Non-absorbed S-RSM4 phage particles. Virions were considered unbound when intact capsids with visible tails and open tail fibers were observed in proximity to a host cell but without direct pilus contact (black arrows). The right-hand panel shows an additional example of an intact virion with clearly visible tail fibers, not adjacent to a cell, included here for illustration purposes only. The same morphological criteria were applied consistently across S-CAM7, S-RSM4, and S-PM2.

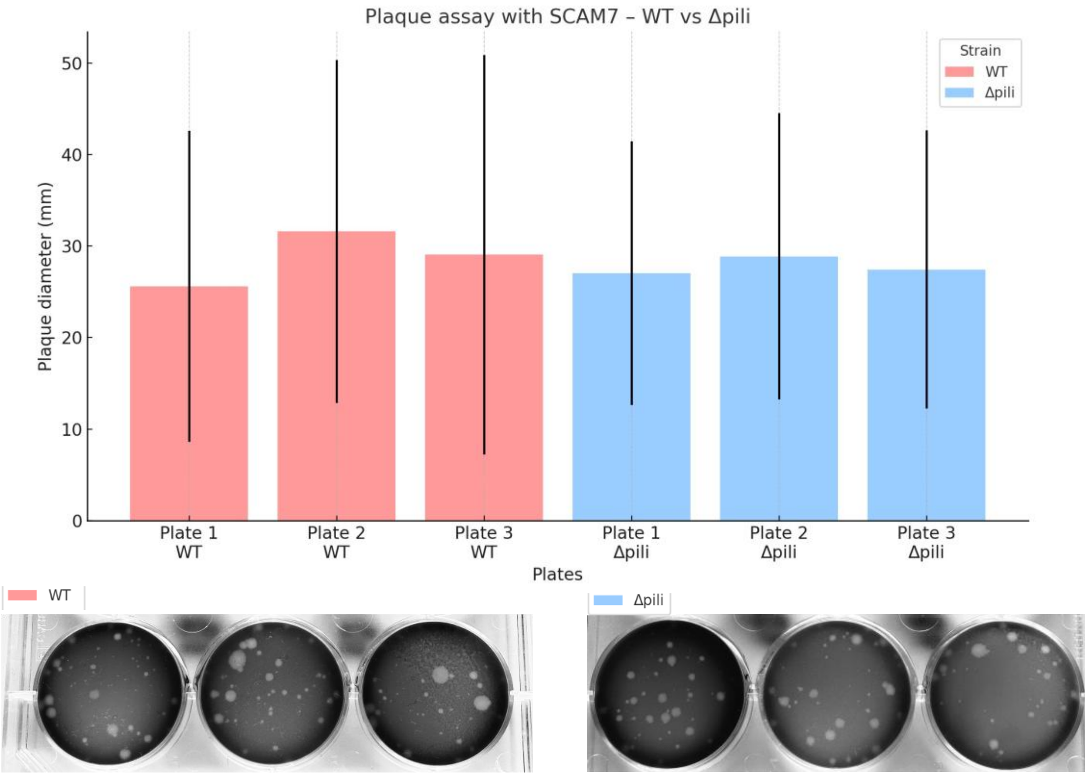


**Supplementary Figure S3. Comparison of plaque sizes in WT and Δpili *Synechococcus* sp. 7803 infected with cyanophage S-CAM7. (A)** Quantification of plaque diameters measured using ImageJ from three replicate plates per strain. **(B)** Representative plaque images for WT (left) and Δpili (right) strains. Cultures were infected with 10 PFU $mL^{-1}$ of S-CAM7 and incubated for 7 days under continuous light. Cell concentrations were determined by flow cytometry: WT = $1.4 \times 10^8$ cells $mL^{-1}$; Δpili = $1.0 \times 10^8$ cells $mL^{-1}$. Plaque diameters were comparable between both strains. This suggests that plaque assays alone may not discriminate between pili-dependent and pili-independent infection pathways. Additionally, type IV pili are structurally fragile and detach during centrifugation or handling steps involved during plaque assay preparation. Furthermore, pili production by the wild type strain within the sloppy agar plates is unknown.

### Supplementary Note 1. Smoluchowski adsorption model for piliated versus non-piliated cells

To estimate how type IV pili change host–phage encounter rates, we used a simple adsorption framework based on the Smoluchowski model. In this model, the adsorption constant depends on the effective target size of the cell that a diffusing phage can collide with. For a non-piliated *Synechococcus* sp. WH7803 cell, the radius is ~0.5 μm (cell diameter 1 μm). When type IV pili are present, they can extend up to 10 μm into the surrounding medium. If we treat this extension as an encounter "envelope", where a collision with a pilus counts as a collision with the cell, the effective radius increases from 0.5 μm to 10 μm. Because the adsorption constant scales linearly with this radius, the presence of pili can increase the theoretical adsorption rate by roughly 20-fold.

This number should be seen as an upper bound rather than an exact prediction. The actual increase will depend on how many pili are expressed, how they are oriented, how dynamic they are (extension/retraction), and whether collisions with pili have the same probability of leading to irreversible adsorption as collisions on the cell surface.

In other words, pili act as **encounter amplifiers**: even if not all collisions result in binding, their extension into the surrounding seawater substantially increases the chances of a phage encountering its host.
